# Deep Neural Networks Predict MHC-I Epitope Presentation and Transfer Learn Neoepitope Immunogenicity

**DOI:** 10.1101/2022.08.29.505690

**Authors:** Benjamin Alexander Albert, Yunxiao Yang, Xiaoshan M. Shao, Dipika Singh, Kellie N. Smith, Valsamo Anagnostou, Rachel Karchin

## Abstract

Identifying neoepitopes that elicit an adaptive immune response is a major bottleneck to developing personalized cancer vaccines. Experimental validation of candidate neoepitopes is extremely resource intensive, and the vast majority of candidates are non-immunogenic, making their identification a needle-in-a-haystack problem. To address this challenge, we present computational methods for predicting MHC-I epitopes and identifying immunogenic neoepitopes with improved precision. The BigMHC method comprises an ensemble of seven pan-allelic deep neural networks trained on peptide-MHC eluted ligand data from mass spectrometry assays and transfer learned on data from assays of antigen-specific immune response. Compared with four state-of-the-art classifiers, BigMHC significantly improves the prediction of epitope presentation on a test set of 45,409 MHC ligands among 900,592 random negatives (AUROC=0.9733, AUPRC=0.8779). After transfer learning on immunogenicity data, BigMHC yields significantly higher precision than seven state-of-the-art models in identifying immunogenic neoepitopes, making BigMHC effective in clinical settings. All data and code are freely available at https://github.com/KarchinLab/bigmhc.

## Introduction

Class I major histocompatibility complex (MHC) plays a crucial role in vertebrate adaptive immunity. The MHC region is highly polymorphic and comprises thousands of known alleles, each encoding a molecule with varying ligand specificities. Identifying non-self-antigens that are presented by a patient’s MHC molecules and elicit strong immune responses may yield precise immunotherapies^1^. Tumor-specific antigens, called neoantigens, and their antigenic determinants, called neoepitopes, are valuable targets for personalized cancer immunotherapies. However, identifying neoepitopes that elicit an antigen-specific immune response is a needle-in-a-haystack problem; the number of non-immunogenic candidates far surpasses the few immunogenic ones. Because experimental validation of immunogenicity is extremely resource intensive, it is critical that the top neoepitope predictions are immunogenic. To address this challenge, we present a deep neural network ensemble called BigMHC for predicting immunogenic neoepitopes with improved precision. Before discussing the method, we first introduce the background.

Intracellular proteins are degraded by proteasomes, after which the resulting peptides may be carried by transporters associated with antigen processing (TAP) molecules to the endoplasmic reticulum (ER). Within the ER, MHC Class I molecules may bind peptides to form a peptide-MHC (pMHC) complex, which may be presented at the cell surface for T-cell receptor (TCR) recognition and subsequent CD8+ T-cell expansion. The set of peptides in each stage is a superset of the following; in other words, given an MHC molecule *M*, then *C* ⊃ *A* ⊃ *B* ⊃ *P* ⊃ *R* ⊃ *T* ⊃ *S*, where:

- *C* is the set of peptides derived from proteasomal cleavage.
- *A* is the set of peptides transported by TAP molecules to the ER.
- *B* is the set of peptides that binds to *M* to form a pMHC complex.
- *P* is the set of peptides present on the cell surface.
- *R* is the set of peptides that form TCR–MHC complexes.
- *T* is the set of peptides that elicits CD8+ T-cell clonal expansion.
- *S* is the set of peptides that elicit a clinically observable antigen-specific immune response.

Some prior works have explicitly incorporated set *C* by modelling proteasome cleavage^2–4^ and set *A* by estimating TAP transport efficiency^4^. Classifiers of set *B* train on in vitro binding affinity (BA) assay data^1,2, 4–11^; however, BA data does not capture the endogenous processes that yield sets *C* and *A*, so BA data capture a strict superset of set *B*. BA data may be qualitative readings or quantitative half-maximal inhibitory concentration (IC_50_) data. Qualitative readings were mapped to IC_50_ values^12^, and the domain of IC_50_ measurements was scaled such that the weakest binding affinity of interest, 5 × 10^5^ nmol/L, was mapped to 0 accordingly: *f*(*IC*_50_) = max(0,1 − log_5×10^5^_(*IC*_50_)).

Mass spectrometry (MS) data, referred to as eluted ligands (EL), are naturally presented MHC ligands; MS data implicitly captures sets *C*, *A*, and *B* while explicitly representing set *P*. EL data provide positive training examples and random pMHC data are generated for negative training examples. Some models^1,2,6, 9–11^ train on both BA and EL data, whereas other models^3,13^, including BigMHC, do not train on BA. To classify set *R*, a recent method^14^ incorporated CDR3β sequences from the TCR to predict the binding affinity between TCR and pMHC. Although TCR information may be useful for predicting immunogenicity, most current datasets do not include such data. The data for classes *R*, *T*, and *S* were also very limited, making it difficult to train classifiers directly for these sets. Prior classifiers of set *T*^15,16^ incorporate the predictions from classifiers of sets *B* and *P*. To the best of our knowledge, there are no predictors of set *S*.

We briefly overview seven state-of-the-art methods to which we compare with the proposed BigMHC (bɪg mæk) method. NetMHCpan-4.1^6^ predicts sets *B* and *P* using an ensemble of 100 single-layer neural networks. This group introduced the idea of a pan-allelic network^7,8^, which consumes a peptide and a short representation of an MHC allele of interest, thereby allowing a single model to generalize across MHC alleles. NetMHCpan-4.1, like many prior models, predicts raw scores in the range [0,1] in addition to a percent rank output in the range [0,100] which normalizes the score to a given allele. MHCflurry-2.0^2^ is a pan-allelic method that predicts sets *B* and *P* using an ensemble of multilayer feed-forward networks, convolutional networks, and logistic regression. MHCflurry-2.0 optionally consumes N-terminal and C-terminal flanking regions to explicitly model set *C*. TransPHLA^11^ is a pan-allelic method that predicts set *P*, using a transformer-based model. MHCnuggets^1^ predicts set *B*, using allele-specific Long Short-Term Memory (LSTM) networks. HLAthena^3^ has pan-allele models that predict set *P* with single-layer neural networks and optionally consume transcript abundance and peptide flanking sequences. MixMHCpred^9,10,13^ predicts set *P*, using a mixture model and position weight matrices (PWM) to extract epitope motifs. PRIME^13,15^ is an extension of MixMHCpred to predict set *T* and was designed to infer the mechanisms of TCR recognition of pMHC complexes.

Using the procedure illustrated in Fig. 1, we developed two BigMHC models: BigMHC EL and BigMHC IM. To predict set *P*, BigMHC EL trains on eluted ligand mass spectrometry data and random negatives. Then, using BigMHC EL as a base model, BigMHC IM transfer learns directly on immunogenicity data to predict set *T*. Because *P* ⊃ *T*, transfer learning narrows the original classification task rather than transferring to an entirely new one. Transfer learning was performed by retraining the final and penultimate fully-connected layers of the base model on immunogenicity data.

**Fig. 1.**
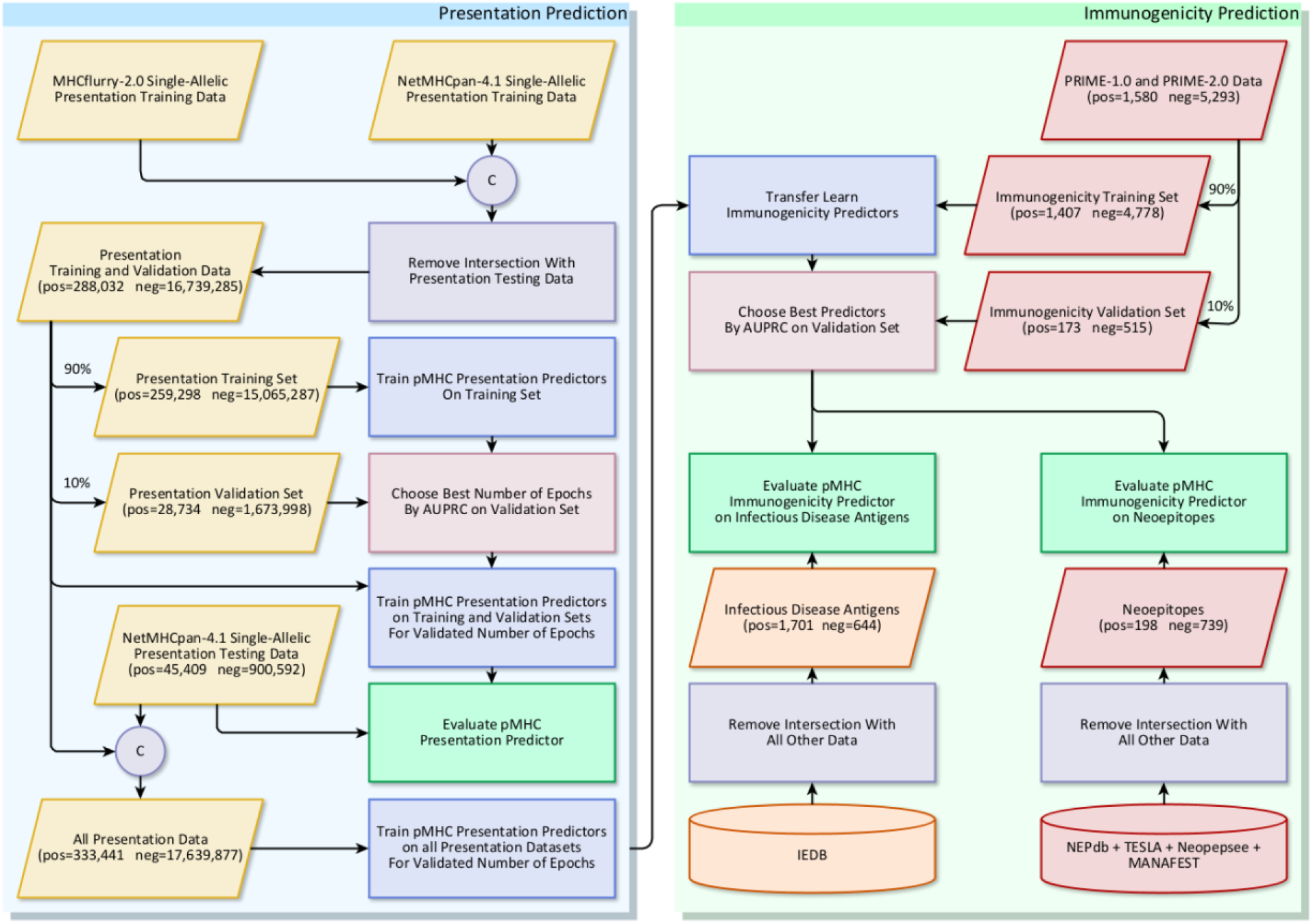
Experimental Procedure. The procedure includes presentation training, immunogenicity transfer learning, and independent evaluation on multiple datasets. The circles labelled ‘C’ indicate dataset concatenation.

Each BigMHC network model (Fig. 2A) is comprised of over 87 million parameters, totaling about 612 million parameters in the ensemble of seven networks. The architecture is designed to capture recurrent patterns via a wide, dense, multilayered, bidirectional LSTM (BiLSTM) and pMHC anchor site binding information via an Anchor Block. The BiLSTM cells are preceded by self-attention modules; these units are equivalent to transformer multi-headed attention modules^17^ where the number of heads is set to 1. Each Wide BiLSTM cell unroll, illustrated in Fig. 2B, consumes the entire MHC representation while recurrently processing the variable-length epitope. Although this imposes a minimum epitope length of 8, few peptides of length 7 or less are presented^3^. The MHC representations are novel pseudosequences generated from multiple sequence alignment; the 30 positions with highest information content are chosen to represent each allele. These positions are one-hot encoded based on the residues present at the given position, with probabilities of occurrence illustrated in Fig. 2C. The Anchor Block consumes the MHC pseudosequence along with the first and last four residues of the peptide to focus on the anchor site residues. The Anchor Block is comprised of two dense^18^ linear layers with tanh activations, followed by Dropout^19^ units with a probability of 0.5. The outputs of the BiLSTM and the Anchor Block are concatenated before being consumed by a Pre-Attention Block, which is also comprised of two dense linear layers with tanh activations, proceeded by Dropout with a probability of 0.5. The output is projected to the same size as the MHC one-hot encoding and passed through tanh activation to attend to the MHC encoding. This attention vector can then be superimposed onto a three-dimensional structure of an MHC allele of interest to identify important amino acid residue positions for a given pMHC, as illustrated in Extended Data Fig. 1 and Extended Data Fig. 2. Moreover, because the final output of the model is a linear combination of the MHC one-hot encoding, the scalar output is interpretable, each MHC position is assigned a weight that contributes in favor of, or against, presentation, and their sum is the model output prior to sigmoid activation.

**Fig. 2.**
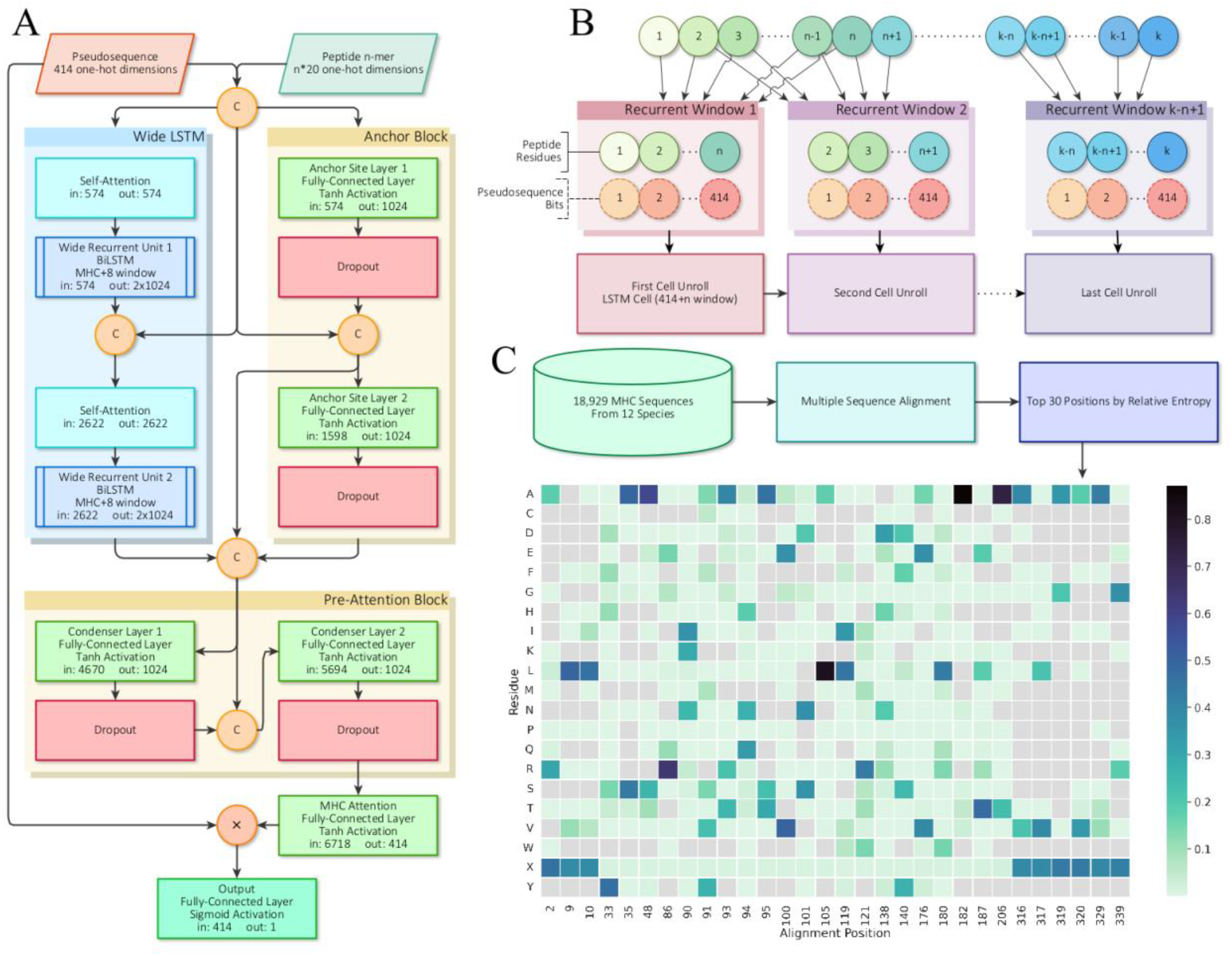
BigMHC network architecture and pseudosequence composition. **A** The BigMHC deep neural network architecture is illustrated, where the BigMHC ensemble is comprised of seven such networks. Pseudosequences and peptides are one-hot encoded prior to feeding them into the model. The circles labelled ‘C’ indicate concatenation and the circle labeled ‘×’ denotes element-wise multiplication. The Anchor Block consists of two densely-connected layers that each receive the first and last four peptide residues along with the MHC encoding. The self-attention modules are single-headed attention units, which is analogous to setting the number of heads of a standard multi-headed Transformer attention module to 1. Prior to the final sigmoid activation, the output of the model is a weighted sum of the MHC pseudosequence onehot encoding; the weights are referred to as attention. Because all connections except internal BiLSTM cell connections are dense, data are not bottlenecked until the MHC attention node maps the pre-attention block output to a tensor of the same shape as the onehot-encoded MHC pseudosequences. **B** A wide LSTM is illustrated. Each cell unroll processes the entire MHC pseudosequence but only a fixed-length window of the peptide. Where a canonical LSTM uses a window of length 1, BigMHC uses a window of length 8 to capitalize on the minimum pMHC peptide length. **C** The pseudosequence residue probability per alignment position. Note that not all residues are present for each position, as indicated by gray cells, so the one-hot encoding uses a ragged array, encoding only the residues present at a given position.

We compared the features of BigMHC with those of seven state-of-the-art methods, as shown in Table 1. The methods are listed left-to-right in descending order of publication year. The training and prediction data modalities of each method were as follows: binding affinity (BA), eluted ligand presentation (EL), and immunogenicity (IM). We also included information on whether the models are retrainable, open-source, offer GPU acceleration, minimum and maximum peptide length, allow additional context such as flanking sequences or gene expression data, webserver availability, and peptide amino acid restrictions.

**Table 1.** Method and feature comparison of BigMHC and prior works. Cells with ‘Y’ indicate that the method has the given feature. Training rows indicate the type of data on which models trained, whereas Prediction rows indicate what type of peptides the model explicitly predicts. BA indicates binding affinity, EL indicates eluted ligand (presentation), and IM indicates immunogenicity. Models that are provided with executables or source code for retraining on new data are considered retrainable. Pan-allele methods are ones that encode the MHC sequence to generalize predictions across alleles rather than employing multiple allele-specific models. Optional extra context refers to any optional input, such as N-terminal and C-terminal flanking sequences or gene expression data. Models that can consume wild-type amino acids, such as ‘X’, are indicated as such in the final row.

|  |  | BigMHC<br>(Proposed) | PRIME-2.0 | MixMHCpred-2.2 | TransPhLA | PRIME-1.0 | MixMHCpred-2.1 | MHCflurry-2.0 | NetMHCpan-4.1 | MHCnuggets-2.4.0 | HLAthena |
| --- | --- | --- | --- | --- | --- | --- | --- | --- | --- | --- | --- |
| Publication Year |  | 2023 | 2023 | 2023 | 2022 | 2021 | 2020 | 2020 | 2020 | 2020 | 2020 |
| Training | BA |  |  |  | Y |  | Y | Y | Y | Y |  |
|  | EL | Y |  | Y | Y |  | Y | Y | Y | Y | Y |
|  | IM | Y | Y |  |  | Y |  |  |  |  |  |
| Prediction | BA |  |  |  |  |  |  | Y | Y | Y |  |
|  | EL | Y |  | Y | Y |  | Y | Y | Y | Y | Y |
|  | IM | Y | Y |  |  | Y |  |  |  |  |  |
| Optional Extra Context |  |  |  |  |  |  |  | Y |  |  | Y |
| Retrainable |  | Y |  |  |  |  |  | Y |  | Y | Y |
| Transfer Learning |  | Y |  |  |  |  |  |  |  |  |  |
| Open Source |  | Y | Y | Y | Y | Y | Y | Y |  | Y | Y |
| Pan-Allele |  | Y |  |  | Y |  |  | Y | Y |  | Y |
| Optional Single-GPU |  | Y |  |  | Y |  |  | Y |  | Y |  |
| Optional Multi-GPU |  | Y |  |  |  |  |  |  |  |  |  |
| Has Webserver |  |  | Y | Y | Y | Y | Y |  | Y |  | Y |
| Min peptide length |  | 8 | 8 | 8 | 8 | 8 | 8 | 5 | 8 | None | 8 |
| Max peptide length |  | None | 14 | 14 | 15 | 14 | 14 | 15 | None | None | 11 |
| Allows WT AA 'X' |  | Y |  |  | Y |  |  |  | Y | Y |  |

## Results

### Epitope Presentation Prediction

BigMHC, NetMHCpan-4.1, TransPHLA, MixMHCpred-2.1 and MHCnuggets 2.4.0 are first evaluated on a set of 45,409 eluted ligands (set *P*) and 900,592 random decoys serving as negatives. This dataset is the same as that used to evaluate NetMHCpan-4.1^6^, but with 140 deduplicated instances. Some prior methods^2,3,13,15^ could not be evaluated on this dataset as they trained on the eluted ligands or do not predict set *P*.

The results, illustrated in Fig. 3A, suggest that BigMHC improves EL predictive capability, reaching 0.9733 mean AUROC and 0.8779 mean AUPRC when stratifying by MHC. The best prior method was NetMHCpan-4.1 Ranks, with 0.9496 mean AUROC and 0.8329 mean AUPRC. The distributions of AUROC and AUPRC across HLA loci are illustrated for each classifier in Fig. 3B. BigMHC demonstrates strong performance across HLA loci, whereas the performance of other methods degrades slightly on HLA-A, and particularly HLA-C. The median PPVn across alleles, as previously calculated^6^, for each method are: BigMHC (0.8617), NetMHCpan-4.1 Ranks and Scores (0.8279), MixMHCpred-2.1 Ranks (0.7907), MixMHCpred-2.1 Scores (0.7898), TransPHLA (0.6839), and MHCnuggets-2.4.0 (0.6507).

**Fig. 3.**
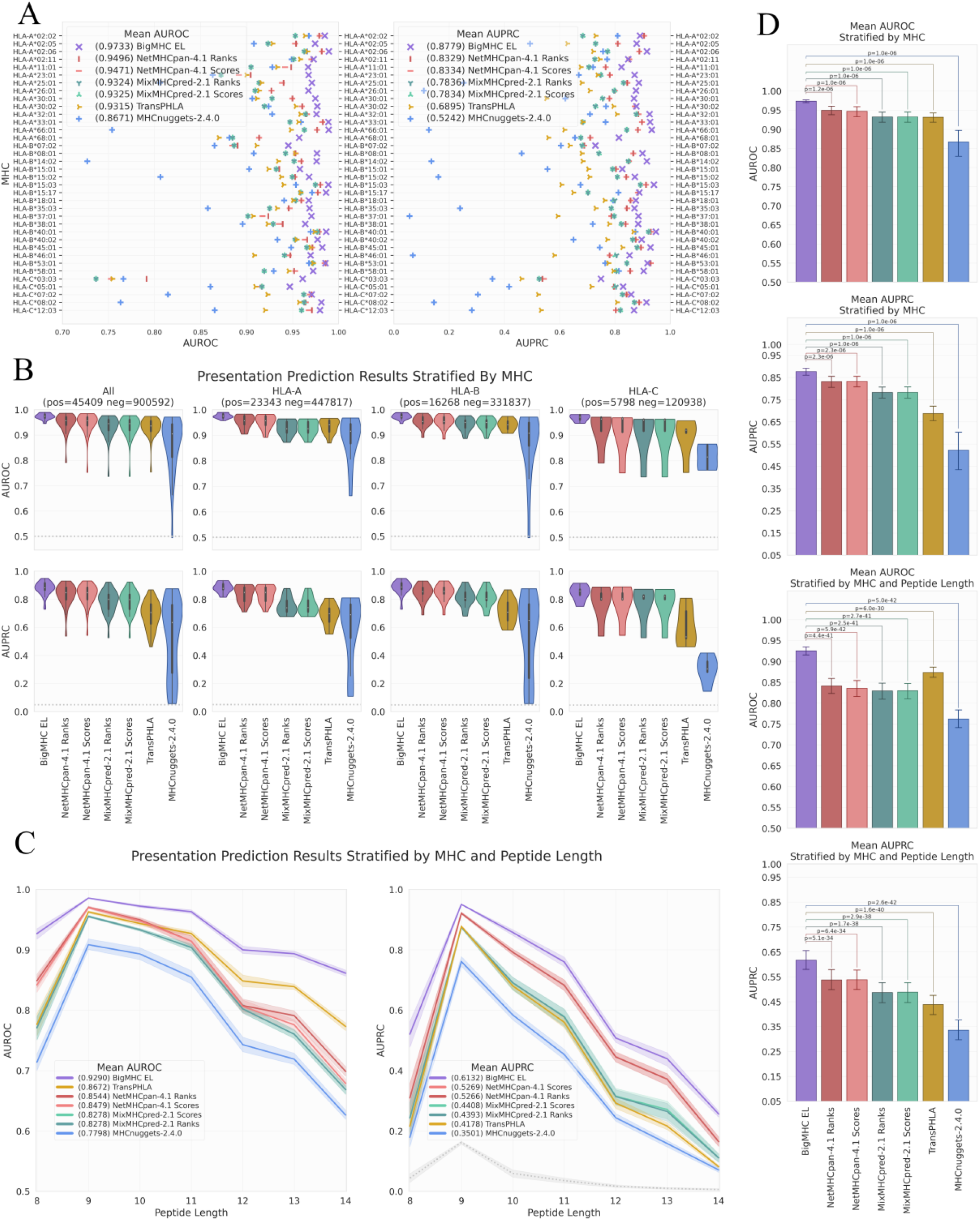
Eluted Ligand prediction results. **A** Area Under the Receiver Operating Characteristic (AUROC) and Area Under the Precision-Recall Curve (AUPRC) for each allele in the EL testing dataset. **B** AUROC and AUPRC violin plots with embedded box-and-whisker plots stratified by allele and grouped by MHC locus. **C** AUROC and AUPRC per peptide allele length with 95% confidence interval by MHC stratification. Baseline (random) classifier performance is 0.5 for AUROC and illustrated in gray for AUPRC. **D** Mean AUROC and AUPRC and 95% confidence interval per stratification with two-tailed Wilcoxon signed-rank test adjusted p-values across methods.

We further stratify by both MHC and peptide length, as illustrated in Fig. 3C. After applying this more granular stratification, BigMHC yields mean AUROC and AUPRC of 0.9290 and 0.6132 respectively. By comparison, NetMHCpan-4.1 yielded mean AUROC and AUPRC of 0.8544 and 0.5266 respectively. BigMHC is most effective for peptides of length 9, which are the most common lengths of peptides presented by MHC-I^3,10^. Although the predictive capability of BigMHC decreases as the peptide length increases, it is still superior to that of the compared methods for all peptide lengths. Overall, BigMHC achieves higher AUROC and AUPRC than these prior methods across both types of stratifications. The two-tailed Wilcoxon signed-rank tests illustrated in Fig. 3D suggest that the BigMHC improvements are statistically significant (adjusted p < 0.05) after Bonferroni correction across the number of compared predictors.

### Immunogenicity Prediction

The vast majority of neoepitopes are not immunogenic. Furthermore, experimental validation of immunogenicity currently is not high-throughput, so it is necessary to select a short list of candidate neoepitopes that can be validated per patient in a clinical setting. It is therefore critical that predictors of immunogenic neoepitopes have high precision among their most highly ranked outputs, as only the top predictions are used in practice. To measure this precision, it is common to evaluate positive predictive value (PPV) among the top n outputs (PPVn)^1,2,6^. To calculate PPVn, the pMHCs are first sorted by a predictor’s output. Then, PPVn is the fraction of the top n pMHCs that are actually immunogenic.

Evaluation of immunogenicity prediction is conducted on two independent datasets: one comprised of neoepitopes and the other comprised of infectious disease antigens. The precision of predicting immunogenic neoepitopes is shown in Fig. 4A; we plot PPVn against all choices of n such that a perfect predictor yields a PPVn of 1. This shows that the top nine predictions are all immunogenic, and as the number of predictions increases, the fraction of predictions that are actually immunogenic remains well above the PPVn of prior methods for all n. To summarize this PPVn curve, the mean PPVn is plotted with 95% confidence interval whiskers in Fig. 4C, showing that BigMHC IM achieves a mean PPVn of 0.4375 (95% CI: [0.4108, 0.4642]), significantly improving over the best prior method, HLAthena Ranks, which achieves a mean PPVn of 0.2638 (95% CI: [0.2572, 0.2705]). Additionally, these data demonstrate the utility of transfer learning to the immunogenicity domain as BigMHC IM significantly outperforms BigMHC EL, which achieves mean PPVn of 0.2704 (95% CI: [0.2632, 0.2776]). A third BigMHC curve is plotted, called BigMHC ELIM, for which we use BigMHC IM predictions on HLA-A and HLA-B peptides and BigMHC EL predictions for HLA-C peptides. Because there were very few HLA-C instances on which to transfer learn, we hypothesized that BigMHC IM may struggle with HLA-C prediction. BigMHC ELIM improved neoepitope immunogenicity AUROC and AUPRC, but did not improve on the infectious disease dataset; this is likely due to the neoepitope dataset being enriched in negative HLA-C peptides compared to the infectious disease dataset, as seen in Extended Data Fig. 5. BigMHC IM and BigMHC ELIM mean PPVn is not significantly different for neoepitope immunogenicity and slightly degrades (adjusted p < 0.05) for infectious disease immunogenicity as determined by two-tailed Wilcoxon signed-rank test with Bonferroni correction.

**Fig. 4.**
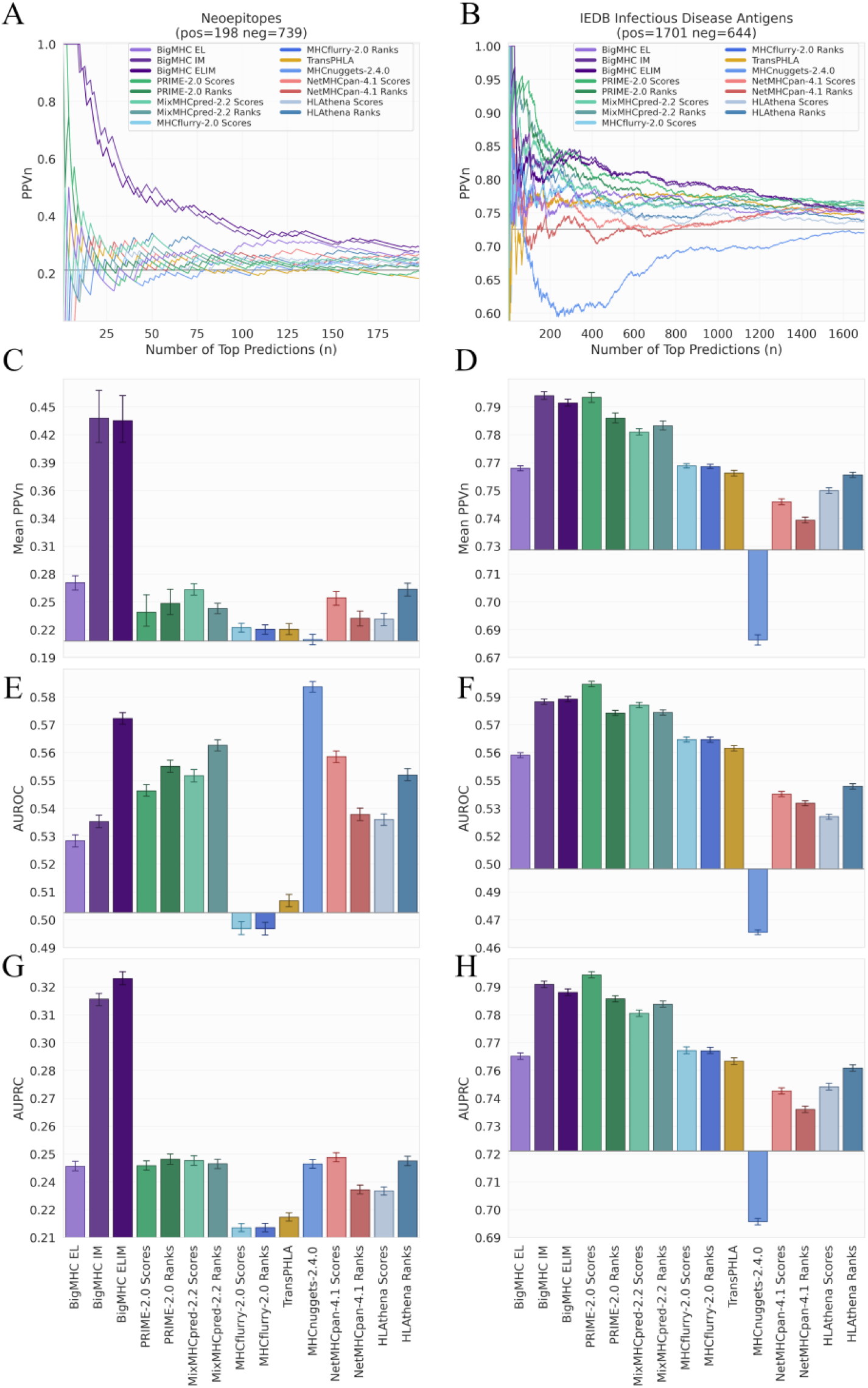
Performance of immunogenicity predictions for all methods. PPVn is calculated for each method as the fraction of (**A**) neoepitopes or (**B**) infectious disease antigens that are immunogenic within the top n predictions. The mean PPVn and 95% confidence interval whiskers are reported (**C-D**), summarizing the PPVn curves for all valid choices of n. The baseline PPVn, representing a random classifier, is illustrated as a horizontal line at 0.2113 for neoepitopes and 0.7254 for infectious disease antigens. AUROC (**E-F)** and AUPRC (**G-H**) of all methods with 95% bootstrap confidence intervals from 1000 iterations on both immunogenicity test sets.

Precision curves and the corresponding mean PPVn for the infectious disease antigen dataset is illustrated in Fig. 4B and Fig. 4D respectively. BigMHC IM achieves a mean PPVn of 0.7999 (95% CI: [0.7980, 0.8018]). The best prior method, PRIME-2.0 Scores, achieves a mean PPVn of 0.7991 (95% CI: [0.7967, 0.8015]). The two-tailed Wilcoxon signed-rank test suggests that the difference between BigMHC IM and PRIME-2.0 Scores is asymmetric about zero (adjusted p < 0.05), but because BigMHC IM just barely improves over PRIME-2.0 precision, we consider these two methods comparable for infectious disease immunogenicity prediction precision.

In addition to precision, we also report AUROC and AUPRC for each dataset along with 1000-fold bootstrapped 95% confidence intervals. The AUROC scores for the neoepitope and infectious disease immunogenicity datasets are reported in Fig. 4E and Fig. 4F respectively, and the AUPRC scores are in Fig. 4G and Fig. 4H respectively. MHCnuggets-2.4.0 achieved the highest AUROC on the neoepitope dataset at 0.5852 (95% CI: [0.5833, 0.5862]), and BigMHC ELIM achieved the next best AUROC at 0.5736 (95% CI: [0.5721, 0.5750]). BigMHC ELIM significantly outperformed all prior methods on neoepitope AUPRC, reaching a mean AUPRC of 0.3234 (95% CI: [0.3216, 0.3253]), whereas the best prior method, NetMHCpan-4.1 Scores, yielded a mean AUPRC of 0.2462 (95% CI: [0.2441, 0.2483]). BigMHC IM yielded AUROC of 0.5348 (95% CI: [0.5332, 0.5363]) and AUPRC of 0.3147 (95% CI: [0.3129, 0.3165]) on the neoepitope dataset, whereas BigMHC EL yielded mean AUROC of 0.5264 (95% CI: [0.5249, 0.5280]) and AUPRC of 0.2415 (95% CI: [0.2401, 0.2428]), further demonstrating significant improvement after transfer learning.

On the infectious disease antigen dataset, PRIME-2.0 Scores achieved the best AUROC and AUPRC, reaching 0.5953 (95% CI: [0.5940, 0.5966]) and 0.7905 (95% CI: [0.7893, 0.7916]) respectively. BigMHC ELIM achieved the next best AUROC at 0.5876 (95% CI: [0.5863, 0.5890]) and BigMHC IM achieved the next best AUPRC at 0.7869 (95% CI: [0.7856, 0.7882]), though both BigMHC IM and BigMHC ELIM yielded similar AUROC and AUPRC on this dataset. As with the neoepitope dataset, both BigMHC IM and BigMHC ELIM improved AUROC and AUPRC over BigMHC EL. The best AUROC and AUPRC for both immunogenicity datasets are statistically higher (adjusted p < 0.05) than the next best as suggested by two-tailed Wilcoxon signed-rank tests with Bonferroni corrections.

### MHC Attention

The BigMHC network architecture offers a unique attention mechanism whereby prior to sigmoidal activation, the scalar output of the network is a linear combination of the input MHC encoding. Hence, we were able to visualize interpretable attention, the amino acid residue positions important for classification, in the form of a heatmap overlay on a modelled three-dimensional structure of an MHC molecule of interest. The mean attention for each pseudosequence position per allele in the EL evaluation dataset is illustrated in Extended Data Fig. 1A. The MHC molecules from each HLA locus that yielded the highest AUPRC are visualized with attention coloring in Extended Data Fig. 1B. The EL training set attention values are visualized in Extended Data Fig. 2. The proposed MHC pseudosequences are comprised of the top 30 aligned positions from a cross-species alignment of 18,929 MHC-I sequences by information content; the most important are those that are in the binding groove. For certain alleles, however, some transmembrane and intracellular residues strongly contribute to EL prediction, such as position 320 for HLA-C*07:02 and 329 for many HLA-B alleles. This suggests that the NetMHCpan pseudosequences, which capture positions only nearest to the peptide, may lose information valuable for predicting pMHC presentation. Importantly, this affects all the referenced pan-allele state-of-the-art methods as they currently adopt NetMHCpan pseudosequence MHC representations.

### Network Architecture Study

We further investigated how the two primary modules of the network architecture affect the BigMHC performance. Specifically, we studied the Wide LSTM architecture and the Anchor Block modules. To perform this investigation, we first ablated the Anchor Block and evaluated this modified architecture. Then, in addition to the Anchor Block ablation, we reverted the Wide LSTM to the canonical implementation and evaluated the resulting model. These two studies suggest that the Wide LSTM and Anchor Block both offer modest gains in performance. We did not ablate the LSTM because the Anchor Block alone is unable to differentiate between peptides of different lengths but with the same first and last four residues. Therefore, an ablated model with the Anchor Block alone would be unable to correctly map the input domain to the target outputs.

#### Anchor Block

The Anchor Block processes the first four and last four residues of the peptide, which we hypothesized would help BigMHC focus on the anchor site binding residues and encourage learning long-range interactions. We ablated the Anchor Block and, using this modified architecture, reconducted the training and evaluation protocols. When stratifying by allele and peptide length, the Anchor Block improved the EL AUROC by 0.0041 and EL AUPRC by 0.0058, and particularly improved on longer peptides (12-14 amino acid residues). However, these differences were not significant at the 0.05 significance level as determined by the two-tailed Wilcoxon signed-rank test. The Anchor Block improved neoepitope immunogenicity mean PPVn by 0.0061 (p=0.046), AUPRC by 0.0064 (p=1.6 × 10^−17^), but worsened AUROC by 0.0038 (p=9.7 × 10^−10^), as determined by the two-tailed Wilcoxon signed-rank test.

#### Wide LSTM

Where a canonical LSTM implementation^1^ recurrently processes a single amino acid residue per LSTM cell unroll, we increase this window so that the BigMHC Wide LSTM processes 8 amino acid residues per cell unroll, overlapping each window by 7 residues as illustrated in Fig. 1B. In this ablation, we compare BigMHC with the Wide LSTM to BigMHC with the canonical LSTM, and neither of these implementations include the Anchor Block. The Wide LSTM implementation may reduce burden on the LSTM cell memory management at the expense of forcing a minimum peptide length of 8. However, most methods impose this restriction, as seen in Table 1, because most peptides presented by MHC-I are at least 8 amino acids in length^10^, so imposing a minimum peptide length via the Wide LSTM does not significantly limit BigMHC usage. Because the Wide LSTM recurs 7 fewer times than the canonical LSTM, the peptide interactions that it must learn are inherently shorter. The Wide LSTM is also faster, improving execution speed per network on the EL test set by nearly 30%. The canonical LSTM required more memory than the Wide LSTM, likely due to more cell unrolls, so the models of batch size 32768 could not be trained. When stratifying by allele and length, the Wide LSTM improved EL AUPRC by 0.0251 (p=2.5 × 10^−14^), and although the canonical LSTM had higher AUROC by 0.0039, that difference was not significant (p=0.062) at the 0.05 significance level from the two-tailed Wilcoxon signed-rank test. For neoepitope immunogenicity prediction, the Wide LSTM improved AUROC by 0.0024 (p=5.9 × 10^−5^), and although the Wide LSTM had higher mean PPVn by 0.0021 (p=0.41) and higher AUPRC by 0.0011 (p=0.056), the latter two differences are not significant at the 0.05 significance level as determined by the two-tailed Wilcoxon signed-rank test.

## Discussion

We first trained BigMHC to predict peptide presentation (set *P*) because an enormous amount of eluted ligand mass spectrometry data for MHC class I peptide presentation is publicly available, making it feasible to train deep learning models with over 87 million parameters. BigMHC EL achieved the highest predictive capability for set *P*, significantly outperforming the four compared methods across HLA loci and epitope lengths. We further demonstrated several technical findings: training on pMHC binding affinity is unnecessary for predicting pMHC presentation, information content is a useful approach for deriving new MHC pseudosequence representations, and some transmembrane and intracellular MHC positions may be important for presentation prediction.

While the goal of neoepitope prediction is ultimately to predict neoepitopes that induce a clinically observable antigen-specific immune response (set *S*), there is limited immunogenicity data to train deep learning models. To address the data scarcity problem, after initially training the base models on presentation data (set *P*), we applied transfer learning using immunogenicity data (set *T*) to produce BigMHC IM. We evaluated BigMHC IM and seven other methods on two independent datasets: neoepitope immunogenicity and infectious disease antigen immunogenicity. We demonstrated strong precision on both datasets, but particularly outperformed all prior methods on the neoepitope immunogenicity prediction. We suspect that BigMHC IM outperforms other tools on neoepitope PPVn but performs similarly on the infectious disease set because of the composition of the training data used for transfer learning. This data is a mixture of neoepitopes, cancer-testis antigens, and viral antigens. The neoepitopes represent a majority of the examples; out of 6,873 experimentally validated examples, 5,279 are neoepitopes.

However, there are several significant limitations to this study. Firstly, our evaluation of presentation prediction is based on eluted MHC ligands detected by mass spectrometry, but the negative data are random. Hence, positive data are limited to the detection efficiency of mass spectrometry, and negative data are not guaranteed negative. Two alleles, namely HLA-B*07:02 and HLA-C*03:03, yielded slightly lower AUPRC than other alleles; we suspect that differences in allele performance are primarily caused by differing class imbalances across the peptide length distributions. Another limitation is that BigMHC can only operate on MHC class I data, whereas some other methods^1,6^ can predict both MHC-I and MHC-II presentation. Although we compare against state-of-the-art methods, there are many other such tools that are not compared here, and EL evaluation could not include MHCflurry-2.0, MixMHCpred-2.2, HLAthena as their training data included most, or all, of the presented epitopes. The datasets used in this study do not have contextual information such as epitope flanking sequences and gene expression data, which may improve MHCflurry-2.0 and HLAthena results. We note that we could not compare performance in a leave-one-allele-out cross-validation experiment as NetMHCpan-4.1, MixMHCpred-2.1, and TransPHLA are not retrainable, and training BigMHC is computationally expensive. We note that a major limitation of our pseudosequences is that new alleles cannot be added without needing to retrain BigMHC from scratch; adding new alleles likely changes the resulting multiple sequence alignment, thereby affecting all other pseudosequences. We also could not answer the question as to whether BigMHC IM could discriminate between immunogenic neoepitopes and presented non-immunogenic neoepitopes; such an experiment would require pMHCs to be validated both in immunogenicity assays and mass spectrometry assays, and to our knowledge there is no such dataset available. Lastly, although our study emphasizes the importance of achieving high precision of immunogenicity in the top-ranked predictions, all evaluations were retrospective.

Future work will include prospective evaluation of the predictions of BigMHC with neoepitope immunogenicity assays. We are implementing BigMHC in ongoing computational analyses of mutation-associated neoepitope evolution under the selective pressure of immune checkpoint blockade in neoadjuvant clinical trials of patients with non-small cell lung cancer (NCT02259621), mesothelioma (NCT03918252) and gastro-esophageal cancer (NCT03044613).

## Methods

### Datasets

#### Epitope presentation training

Presentation training data spanned 149 alleles and included 288,032 eluted ligands and 16,739,285 negative instances. The training dataset was compiled from the NetMHCpan-4.1^6^ and MHCflurry-2.0^2^ single-allelic (SA) EL datasets. These instances were split into training (positive=259,298 and negative=15,065,287), and validation (positive=28,734 and negative=1,673,998).

#### Epitope presentation evaluation

The presentation evaluation set spanned 36 HLA alleles and was comprised of 45,409 eluted ligands and 900,592 negative data. The test set is the same SA EL dataset as that used in the NetMHCpan-4.1 study but with 140 deduplicated instances. The pMHC instances that existed in both EL training and EL testing were removed from the training dataset.

#### Immunogenicity training

BigMHC IM transfer learned on PRIME-1.0^10^ and PRIME-2.0^13,15,20^ datasets. The original training data consisted of viral antigens, cancer-testis antigens, neoepitopes, and 9-mer peptides randomly sampled from the human proteome to supplement the negative instances. BigMHC transfer learned only on the nonrandom pMHC data, of which 1,580 are positive and 5,293 are negative. The data were split into training (positive=1,407 and negative=4,778), and validation (positive=173 and negative=515).

#### Infectious disease antigen immunogenicity evaluation

Infectious disease antigen immunogenicity prediction was evaluated using data collected from the Immune Epitope Database (IEDB)^21^ on December 19, 2022. The queries to IEDB included linear peptides, T-cell assays, MHC-I restriction, human hosts, and infectious diseases. Data was further processed to allow all prior methods to be evaluated: only peptides of length at least 8 and at most 11 were kept so that HLAthena could be evaluated, peptides with dummy amino acid ‘X’ were removed as many prior methods cannot handle dummy amino acids, and pMHCs with MHC alleles incompatible with MixMHCpred and PRIME were removed. After removing the intersection with all other pMHC data, a total of 1,701 immunogenic and 644 non-immunogenic infectious disease antigens were collected.

#### Neoepitope immunogenicity evaluation

The neoepitope immunogenicity dataset was compiled using NEPdb^22^ downloaded on December 18, 2022, Neopepsee^23^, TESLA^16^, and data collected from 16 cancer patients using the MANAFEST assay^24^. NEPdb is a database of neoepitopes curated from the literature with a semi-automated pipeline, whereas Neopepsee aggregated neoepitopes from two prior sources. TESLA validated immunogenicity of neoepitope predictions from a global consortium. The MANAFEST data was comprised of 167 immunogenic and 672 non-immunogenic neoepitopes. MANAFEST quantifies antigen-specific T-cell clonotype expansion in peptide-stimulated T-cell cultures. After removing the intersection with all other pMHC data, a total of 198 immunogenic and 739 non-immunogenic neoepitopes were collected. As with the infectious disease antigen immunogenicity dataset, the only peptides kept were of length at least 8 and at most 11 so that HLAthena^3^ could be evaluated and peptides with dummy amino acid ‘X’ were removed.

#### MANAFEST data collection

The MANAFEST neoepitope data was collected and processed using an established protocol^24–26^. It was generated from functional experiments of mutation associated neoantigen (MANA)-stimulated autologous T cell cultures for 16 patients with non-small cell lung cancer (NSCLC). Neoantigen-specific T cells were identified in peripheral blood using the MANAFEST assay as previously described^24–26^. For each case, tumor whole exome sequencing data were utilized to determine non-synonymous mutations and mutation-associated neoantigen candidates matched to each patient’s MHC class I haplotypes were computed as previously described^27^. ImmunoSELECT-R software from Personal Genome Diagnostics^25,28^ was used to select putative neoantigens and the neopeptides were synthesized by JPT Peptide Technologies. ImmunoSELECT-R incorporates several tools, including predicted MHC class I affinity from NetMHCpan 3.0^5^, cytoxic T lymphocyte epitope prediction from NetCTLpan^29^, and average gene expression in TCGA NSCLC^25,28^. T cells were isolated from peripheral mononuclear cells (PBMC) for each case by negative selection (EasySep; STEMCELL Technologies) and cultured for 10 days as previously reported^24–26^. TCR Vβ next-generation sequencing utilizing DNA from cultured CD8+ cells was performed by the Johns Hopkins Fest and TCR Immunogenomics Core Facility (FTIC) using the Adaptive Biotechnologies hsTCRB Kit using survey-level sequencing (Adaptive Biotechnologies, WA). Processed data files were analyzed in the publicly available MANAFEST analysis web application (http://www.stat-apps.onc.jhmi.edu/FEST) to define neoantigen-specific T cell clonotypes. Briefly, following data preprocessing, alignment and trimming, productive frequencies of TCR clonotypes were calculated. Neoantigen-specific T cell clonotypes met the following criteria: (1) significant expansion (Fisher’s exact test with Benjamini–Hochberg correction for FDR, p < 0.05) compared to T cells cultured without peptide; (2) significant expansion compared to every other peptide-stimulated culture (FDR < 0.05); (3) an odds ratio greater than 5 compared to all other conditions; (4) at least 30 reads in the positive well; and (5) at least 2 times higher frequency than background clonotypic expansions as detected in the negative control condition^24–26^.

#### Dataset Compositions

The compositions of all datasets are illustrated in Extended Data Fig. 5.

### BigMHC Training

BigMHC was developed using Python 3.9.13, PyTorch 1.13^30^, and CUDA 11.7 on an AMD EPYC 7443P CPU with 256 GB of DDR4 RAM, and four NVIDIA RTX 3090 each with 24 GB of GDDR6X RAM. The training data was split 9:1 into a training set and a validation set. Training used the AdamW optimizer^31,32^ to minimize binary cross entropy loss. We fine-tuned the optimizer learning rate from our initial guess of 1 × 10^−5^ to 5 × 10^−5^ by maximizing AUPRC on the EL validation dataset. The other AdamW hyperparameters were set to their default values: *λ* = 0.01, *β*_1_ = 0.9, *β*_2_ = 0.999, and *∈* = 10^−8^. Seven such BigMHC EL models were trained with varying batch sizes in {2*^k^* ∀ *k* ∈ {9,10, …, 15}}, with the maximum batch size of 32768 occupying all GPU memory. For each batch size, we chose the number of training epochs that maximized AUPRC on the EL validation data up to 50 maximum epochs. For the seven models trained with batch sizes 512, 1024, …, 32768, the best BigMHC EL epochs were respectively: 11, 10, 14, 14, 21, 30, and 46. On the validation data, these models yielded a mean AUROC of 0.9930 and a mean AUPRC of 0.8592. Then, the EL validation set was concatenated with the EL training set and we trained a new set of seven models for the number of epochs that previously maximized AUPRC. This new ensemble was used to evaluate BigMHC EL performance on the EL testing dataset.

After evaluating EL prediction on the testing data, all EL data were merged to train a third set of seven models using the previously optimal number of EL training epochs. This third ensemble is used as the base model for transfer learning immunogenicity. The immunogenicity training data is similarly split 9:1 for training and validation. We optimized both the batch size and number of epochs for transfer learning for each base model by choosing the batch size and epoch number that maximizes AUPRC on the IM validation data. We search all batch sizes in {2*^k^* ∀ *k* ∈ {3,4,5,6,7}} and all epochs up to 100. In order of least to greatest base model batch size, the best BigMHC IM (batch, epoch) numbers are: (16,23), (16,23), (8,15), (64,62), (32,27), (32,31), and (64,54). On the IM validation data, these models yielded a mean AUROC of 0.7767 and a mean AUPRC of 0.5685.

### MHC Pseudosequence

We introduced a new MHC pseudosequence representation. Prior pan-allele methods^2,11^ adopted the NetMHCpan^6^ pseudosequences, which represent the MHC molecule based on residues estimated to be closest to the peptide, or used Kidera factors^3^ to encode the binding pocket residues. By contrast, BigMHC uses multiple sequence alignments to identify positions with high information content. In total, 18,929 MHC-I sequences across 12 species from IPD-IMGT/HLA^33^, IPD-MHC 2.0^34^, and UniProt^35^ (accession numbers: P01899, P01900, P14427, P14426, Q31145, P01901, P01902, P04223, P14428, P01897, Q31151) were aligned using the buildmodel and align2model from the SAM suite^36–38^ version 3.5 with default parameters, yielding 414 aligned residues per sequence. The top 30 positions by information content were identified using makelogo from the SAM suite and were selected to represent the MHC sequences, which can be one-hot encoded with 414 binary variables. These new pseudosequences are provided in our public Git repository.

### Compared Methods

NetMHCpan-4.1^6^ is a popular tool for simultaneously predicting BA and EL. This method consists of an ensemble of 100 single-layer networks, each of which consumes a peptide 9-mer binding core and a subsequence of the MHC molecule. The 9-mer core is extracted by the model, whereas the MHC representation, termed a “pseudosequence,” is a predetermined 34-mer core extracted from the MHC molecule sequence. The 34-mer residues were selected based on the estimated proximity to bound peptides so that only residues within 4 Å were included.

MHCflurry-2.0^2^ is an ensemble of neural networks that predicts BA and EL for MHC-I. BA prediction is the output of a neural network ensemble, where each member is a 2 or 3 layer feed-forward neural network. Then, an antigen processing (AP) convolutional network is trained on a subset of the BA predictions, along with N-terminus and C-terminus flanking regions, to capture antigen processing information that is missed by the BA predictor. EL prediction is the result of logistically regressing BA and AP outputs. The MHC representation was adopted from NetMHCpan pseudosequences and expanded by three residues to differentiate some HLA alleles.

TransPHLA^11^ is a transformer-based model that adopts NetMHCpan pseudosequences for MHC encoding. This model encodes peptides and MHC pseudosequences using the original transformer encoding procedure^17^ before inferring the encodings via ReLU-activated fully-connected layers. TransPHLA was trained on the BA and EL data, although the BA data were binarized as binding or non-binding instances. We found that the final softmax activation of TransPHLA forces many outputs to precisely 0 or 1, which prevents the calculation of metrics, such as AUROC and AUPRC. Therefore, we removed the final softmax activation from TransPHLA to increase model output granularity. Because softmax is monotonic and none of the evaluation metrics rely on arbitrarily thresholding model outputs, removing the final softmax activation does not affect the evaluation metrics used in this study.

MHCnuggets^1^ is comprised of many allele-specific Long Short-Term Memory (LSTM) networks to handle arbitrary-length peptides for MHC-I and MHC-II. Transfer learning was used across the alleles to address data scarcity. MHCnuggets trained on qualitative BA, quantitative BA, and EL data. MHCnuggets trained on up to two orders of magnitude fewer data than the other methods.

MixMHCpred^9,10,13^ is built using positional weight matrices (PWM) to extract epitope motifs for each allele in their training data for peptides of lengths 8 to 14. MixMHCpred-2.1 was used to evaluate EL performance because MixMHCpred-2.2 trained on the EL testing data. PRIME^13,15^ builds off MixMHCpred, training directly on immunogenicity, and was designed to infer the mechanisms of TCR recognition of pMHC complexes. Upon evaluating MixMHCpred versions 2.1 and 2.2 and PRIME versions 1.0 and 2.0, both of the methods’ newest versions offer substantial improvement over their predecessors.

HLAthena^3^ uses three single-layer neural networks trained on mass-spectrometry data to predict presentation on short peptides with length in the range [8,11]. Each of the three networks trained on a separate peptide encoding: one-hot, BLOSUM62, and PMBEC^39^. In addition, the networks consumed peptide-level characteristics, and also amino acid physicochemical properties. The outputs of these networks were used to train logistic regression models that also accounted for proteasomal cleavage, gene expression, and presentation bias. HLAthena also saw performance gains when considering RNA-seq as a proxy for peptide abundance.

## Ethics Declaration

Under a license agreement between Genentech and the Johns Hopkins University, X.M.S., and R.K., and the University are entitled to royalty distributions related to MHCnuggets technology discussed in this publication. This arrangement has been reviewed and approved by the Johns Hopkins University in accordance with its conflict-of-interest policies. V.A has received research funding to her institution from Bristol Myers Squibb, Astra Zeneca, Personal Genome Diagnostics and Delfi Diagnostics in the past 5 years. V.A is an inventor on patent applications (63/276,525, 17/779,936, 16/312,152, 16/341,862, 17/047,006 and 17/598,690) submitted by Johns Hopkins University related to cancer genomic analyses, ctDNA therapeutic response monitoring and immunogenomic features of response to immunotherapy that have been licensed to one or more entities. Under the terms of these license agreements, the University and inventors are entitled to fees and royalty distributions. The remaining authors have declared no conflicts of interest.

## Data Availability

All data except MANAFEST data were collected from publicly available sources^2,6,15,16, 21–23^. All data, including model outputs, are provided in our public Mendeley repository: https://data.mendeley.com/datasets/dvmz6pkzvb.

## Code Availability

All code used in this study is provided in our public GitHub repository: https://github.com/KarchinLab/bigmhc. In addition, the final trained models are provided in this repository.

## Acknowledgements

This work was supported in part by the US National Institutes of Health grant CA121113, the Department of Defense Congressionally Directed Medical Research Programs grant CA190755, and the ECOG-ACRIN Thoracic Malignancies Integrated Translational Science Center grant UG1CA233259.

## Contributions

B.A.A. and R.K. conceived the study and performed the experiments; Y.Y. contributed to 3D visualizations and model ideas; X.M.S. curated the MANAFEST data; D.S. and K.N.S. collected the MANAFEST dataset; B.A.A. and R.K. wrote the draft manuscript; B.A.A., V.A. and R.K. revised the manuscript; R.K. supervised the research.

## Extended Figures

**Extended Data Fig. 1.**
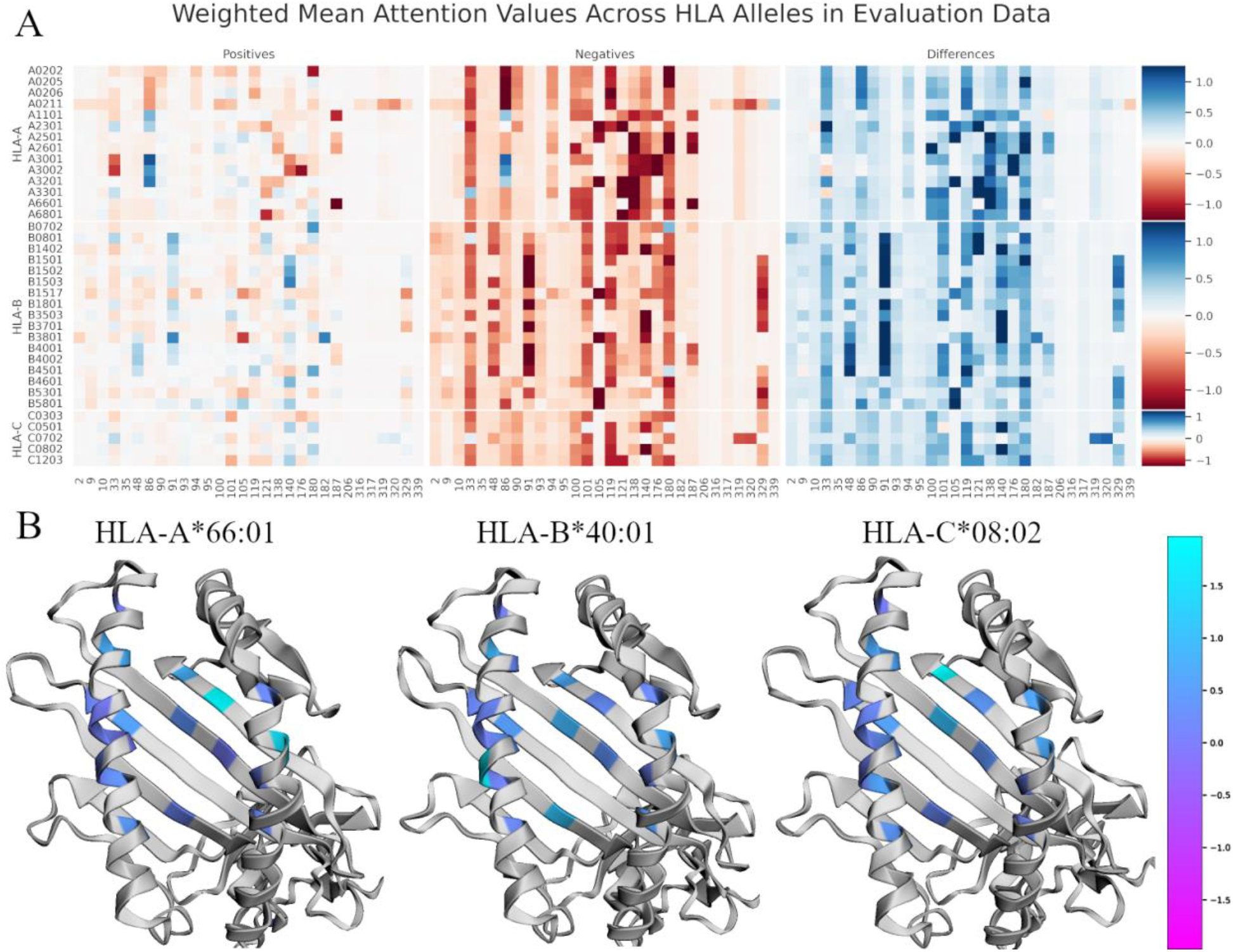
Visualization of BigMHC attention to MHC encodings. **A** Heatmap visualization of the average MHC attention values stratified by MHC allele and presentation class on the EL testing dataset. Darker values indicate MHC positions that are more influential on the final model output. The column of Differences depicts the Negatives values subtracted from the Positives values; thus, darker blue colors are most correctly discriminative whereas darker red attention values in this column highlight erroneous inferences. **B** Overlays of the Differences column from the training dataset on the MHC molecule using py3Dmol^40^. HLA crystal structures are generated using AlphaFold^41^.

**Extended Data Fig. 2.**
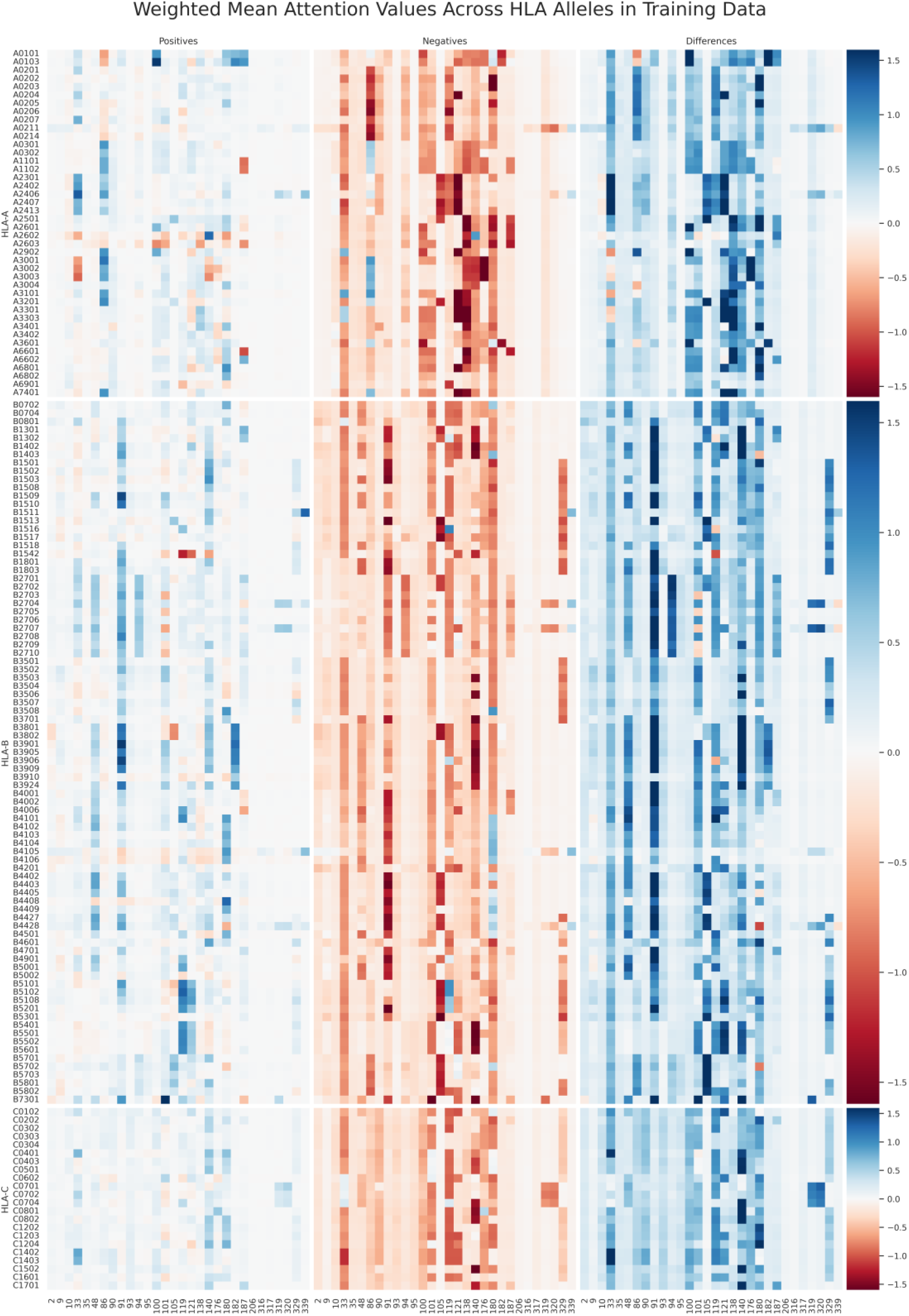
Heatmap visualization of the average MHC attention values stratified by MHC allele and EL target class on the EL training dataset.

**Extended Data Fig. 3.**
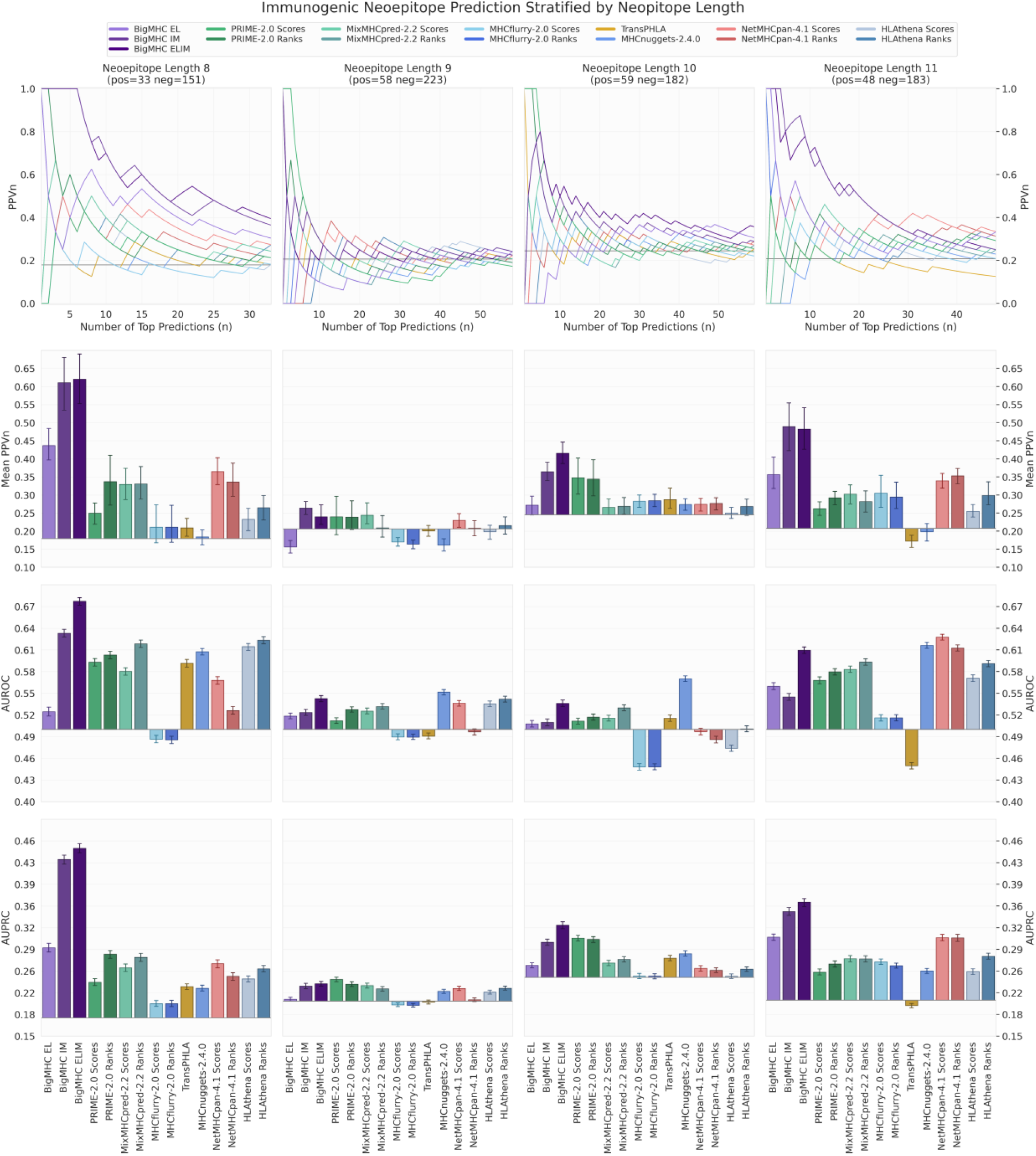
Neoepitope immunogenicity prediction results stratified by neoepitope length.

**Extended Data Fig. 4.**
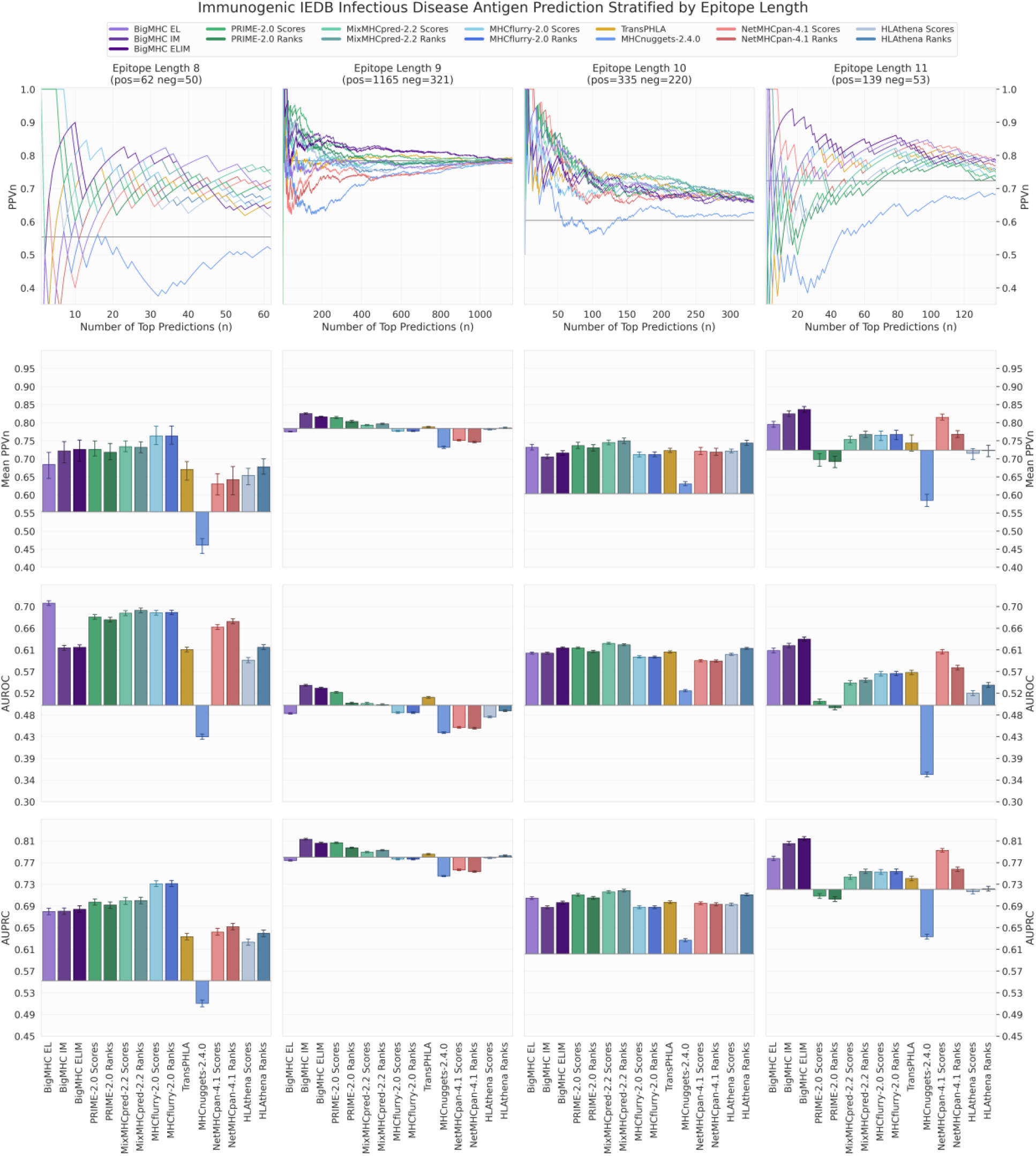
IEDB infectious disease antigen immunogenicity prediction results stratified by epitope length.

**Extended Data Fig. 5.**
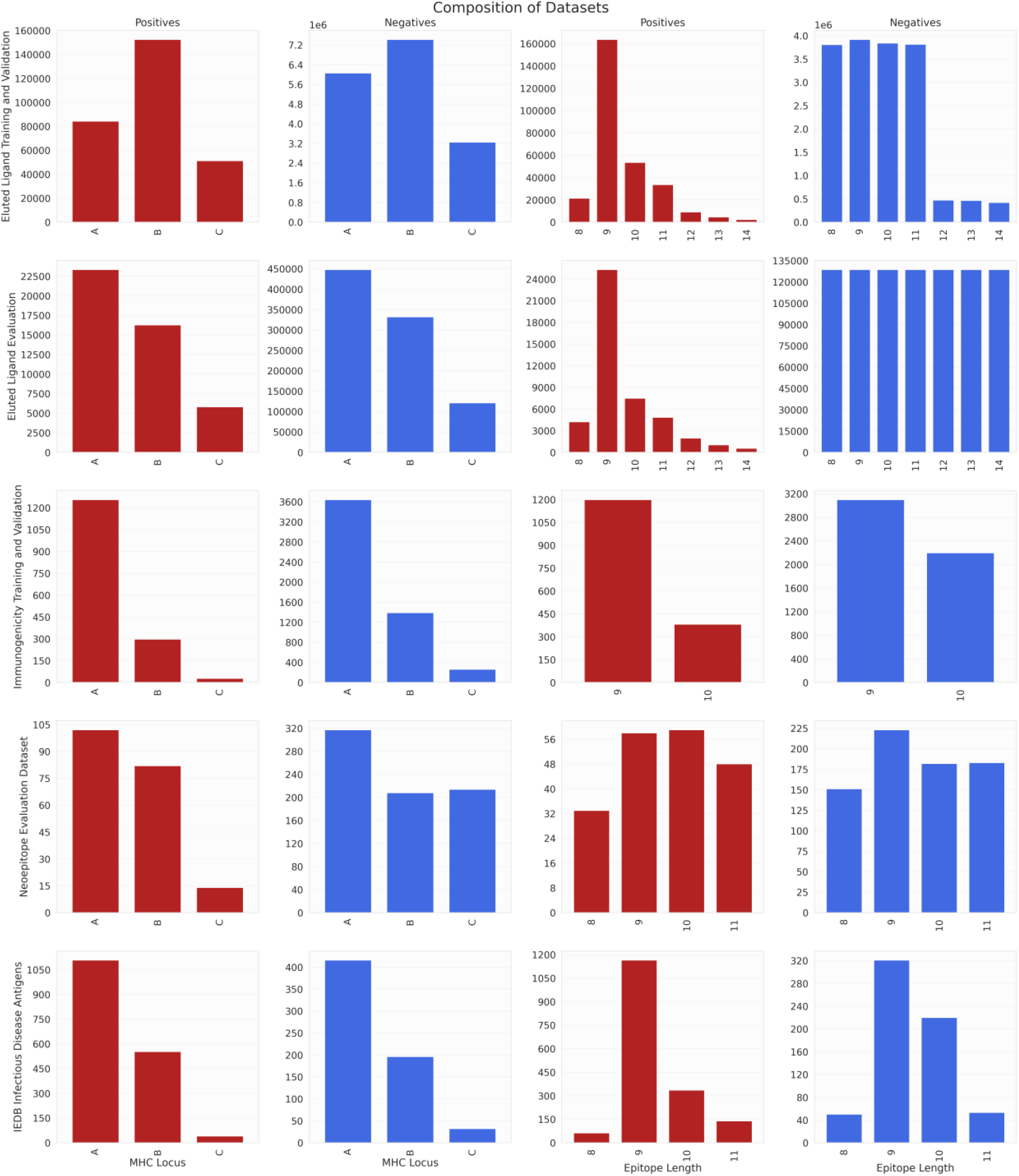
Composition of all training and evaluation datasets. Positive and negative instances were stratified by HLA loci in the first two columns and by epitope length in the latter two columns. Positives in the EL datasets are detected by mass spectrometry, whereas negatives in the EL datasets are decoys. Both positives and negatives in the immunogenicity datasets are experimentally validated by immunogenicity assays.

